# Engineering a soft tumor microenvironment: fibrin-enriched hydrogels promote cancer cell invasion in 3D bioprinted colorectal cancer models

**DOI:** 10.64898/2026.01.08.698319

**Authors:** Melika Parchehbaf Kashani, Jordi Comelles, Pol Barcelona, Núria Torras, Lorena Dieguez, María García-Díaz, Elena Martínez

## Abstract

Accurately modeling the tumor microenvironment is crucial for advancing our understanding of colorectal cancer (CRC) progression and therapeutic response. Three-dimensional (3D) hydrogel-based models that mimic the mechanical properties of native tissue serve as a valuable tool for studying tumor–stromal interactions and tumor invasion in vitro. In this work, we developed a 3D bioprinted CRC model by embedding spheroids and human intestinal fibroblasts (HIFs) within the GelMA–PEGDA and GelMA–PEGDA–Fibrin hydrogels to assess the effect of matrix composition and stiffness on spheroid invasiveness. The addition of fibrin resulted in a softer, more viscoelastic hydrogel that promoted fibroblast migration, elongation, and alignment, facilitating more dynamic tumor–stromal interactions. Moreover, both epithelial-like HT29 and mesenchymal-like SW480 spheroids showed more invasive behavior in GelMA-PEGDA-Fibrin hydrogel. This was reflected in distinct phenotypic responses. HT29 spheroids demonstrated greater growth, irregular morphology, and more interaction with elongated fibroblasts, whereas SW480 spheroids exhibited partial dissociation and disruption with a higher number of dispersed cells in GelMA-PEGDA-Fibrin hydrogel. These findings demonstrate the role of matrix softness in promoting the invasiveness of colorectal cancer.

Overall, our results highlight the potential of fibrin-enriched soft hydrogel to mimic key features of the tumor microenvironment, offering a powerful tool for studying CRC invasion dynamics and supporting future applications in drug screening and personalized medicine.

## Introduction

Colorectal cancer is the third most frequently diagnosed malignancy and the second common cancer-related death worldwide ^1^. The development, progression, and metastatic behavior of colorectal cancer (CRC) are highly influenced by the tumor microenvironment (TME), which is composed of a cellular and a non-cellular compartment. The cellular component includes various cells such as fibroblasts, immune cells, endothelial cells, and pericytes, while the non-cellular component, the extracellular matrix (ECM), is a structured network of fibrous proteins that provides mechanical support and regulates cancer cell invasiveness ^2^. Among the various components of the TME, stromal fibroblasts represent the predominant cell type and play a pivotal role in stromal–tumor crosstalk, which modulates tissue architecture, intercellular signaling, and tumor behavior ^3^. Recent studies have shown that fibroblast migration is crucial in the tumor microenvironment since it remodels the extracellular matrix and creates pathways that promote cancer cell invasion. Migrating fibroblasts direct tumor cells along physical tracks and molecular signals, promoting collective movement and spread. Their activity directly impacts tumor growth, making them essential participants in cancer invasion and prospective therapeutic targets ^4,5^. Moreover, the mechanical properties of TME play a crucial role in the migration and invasion of cancer cells in vivo ^6^. To further investigate the complex tumor-stromal interaction and cancer invasion, the development of physiologically relevant in vitro models is essential. Traditional two-dimensional (2D) in vitro models have been widely used in cancer biology due to their simplicity; however, they fail to mimic the spatial complexity, mechanical cues, and dynamic cell–cell and cell–matrix interactions found in TME ^7,8^. Over the last few decades, hydrogel-based 3D models have become an important tool in cancer research, providing more accurate models of the TME by employing diverse materials and techniques that mimic the native tissue microenvironment ^9^. Among these, the combination of tumor spheroids with hydrogel-based scaffolds is considered an attractive in vitro approach for studying cancer behavior as it combines the complexity of multicellular tumor spheroids with the tunability of hydrogel stiffness representing the TME. In this context, Chen et al have studied gelatin methacryloyl (GelMA), collagen, or hybrid composites to model cancer invasion. They demonstrated that the increase of GelMA crosslinking density, thereby increasing stiffness and reducing pore size, led to a restricted invasive behavior of embedded tumor spheroids ^10^. GelMA is one of the polymers most used in 3D bioprinting, especially in light-based bioprinting. This photocrosslinkable polymer is a biodegradable polymer with cell adhesion motifs and high elasticity, but it is not mechanically stable ^11,12^ In order to increase the printability and long-term stability, it is often combined with inorganic or organic components. In the context of CRC, a 3D CRC model which was bioprinted using GelMA-nanocly hydrogels showed an increase in colorectal cancer stem cells by activating the Wnt/β-catenin signaling pathway. This model supported the formation of spheroids with elevated stemness markers, providing a valuable platform for studying CSC behavior and drug responses ^13^. Polyethylene glycol diacrylate (PEGDA) is another polymer commonly used as bioink. It is a biocompatible hydrogel that provides sufficient mechanical stability. However, it lacks cell adhesion motifs^14,15^ and is used in combination with other components such as thiolated gelatin (GelSH), fibrin, or GelMA to fabricate constructs mimicking tissues like the intestine ^14–16^.

The healthy colon mucosa has been found to be very soft, less than 0.1 kPa as measured from patient’s biopsies ^17,18^. This stiffness significantly increased in CRC tumoral areas and their surrounding stromal tissue but remains very soft in the order of 0.2-0.5 kPa. Obtaining bioprintable hydrogels with that low stiffness range remains challenging, as the low polymer concentration and crosslinking density required to achieve such softness often compromise print fidelity and structural stability during printing. In particular, insufficient mechanical integrity can lead to poor layer definition and deformation during post-print handling, necessitating a careful balance between printability, photopolymerization kinetics, and mechanical compliance. In this work, we explore the combination of a low polymer content GelMA-PEGDA hydrogel and the incorporation of fibrin as interpenetrating network to mimic the colon tumoral mucosa mechanical properties using light-based bioprinting. To mimic tumor microenvironment, HIFs as the most abundant stromal cells and tumor spheroids derived from HT29 and SW480 colorectal cancer cell lines were embedded within the hydrogel bulk. Phenotypic characterization confirmed the more epithelial-like characteristics of HT29 (less invasive) spheroids compared to the more mesenchymal SW480 spheroids (more invasive). The incorporation of the fibrin network within the GelMA-PEGDA hydrogel resulted in a softer matrix that significantly increased the motility of the HIFs, confirming the effect of fibrin on cell migration ^19,20^. Modulating hydrogel stiffness via fibrin incorporation profoundly influenced spheroid growth dynamics, morphology, and invasive behavior over time, as demonstrated by increased growth rates and decreased circularity in HT29 spheroids, alongside a higher number of dissociated cells in SW480 spheroids within the softer GelMA-PEGDA-Fibrin hydrogel. These findings underscore the essential role of hydrogel mechanical properties and composition in directing fibroblast migration, spheroid morphology, and invasiveness, establishing a physiologically relevant platform for studying colorectal cancer progression and advancing preclinical drug screening.

## 2. Experimental section

### 2.1. Gelatin methacryloyl (GelMA) synthesis

GelMA was prepared following the method described previously ^16,21^. Briefly, gelatin solution (10% w/v) was prepared by dissolving porcine skin type A gelatin (Sigma-Aldrich) in phosphate-buffered saline (PBS; pH 7.4) (Gibco) at 50°C while stirring for approximately 2 h. Methacrylic anhydride (MA) (Sigma-Aldrich) at 5% v/v was added at a flow rate of 0.5 mL/min and allowed to react for 1 hour with continuous stirring. After that, the solution was centrifuged at 1200 rpm for 3 minutes, and the reaction was stopped by adding Milli-Q water to the supernatant. The resulting solution was subjected to dialysis using 6-8 kDa molecular weight cut-off membranes (Spectra/por, Spectrumlabs) against Milli-Q water at 40°C, with water replacement every 4 h for 3 days. The pH of the dialyzed products was adjusted to 7.4. Finally, the samples were frozen overnight at −80°C and lyophilized for 4 and 5 days (Freeze Dryer Alpha 1-4 LD Christ). The final product was stored at −20°C until further use. The methacrylation degree was characterized using the 2,4,6-Trinitrobenzene sulfonic acid (TNBS) assay ^22^, resulting in 42 ± 6 % of conjugation.

### 2.2. Printing of cell-free hydrogels

Two different bioink compositions, named GelMA-PEGDA and GelMA-PEGDA-Fibrin, were printed and the formed cell-free hydrogels were characterized (Figure 1A). To prepare the GelMA-PEGDA prepolymer solution, we dissolved 5% w/v of GelMA, 1.25% w/v of PEGDA with a molecular weight of 4000 Da (Polysciences), 0.2% w/v of visible-light lithium phenyl-2,4,6-trimethylbenzoylphosphinate (LAP) photoinitiator (TCI Europe), and 0.025% w/v of tartrazine food dye (Acid Yellow 23, Sigma-Aldrich) at 65°C in Hank’s Balanced Salt (HBSS) solution complemented with 1% Penicillin-Streptomycin (Sigma-Aldrich) as dilution buffer (Sigma-Aldrich) for 2 h under stirring conditions. For the GelMA-PEGDA-Fibrin, 0.25% w/v of fibrinogen (Merck, Life Science) was mixed with the GelMA-PEGDA prepolymer solution and kept at 37°C for 30 min before use.

**Figure 1.**
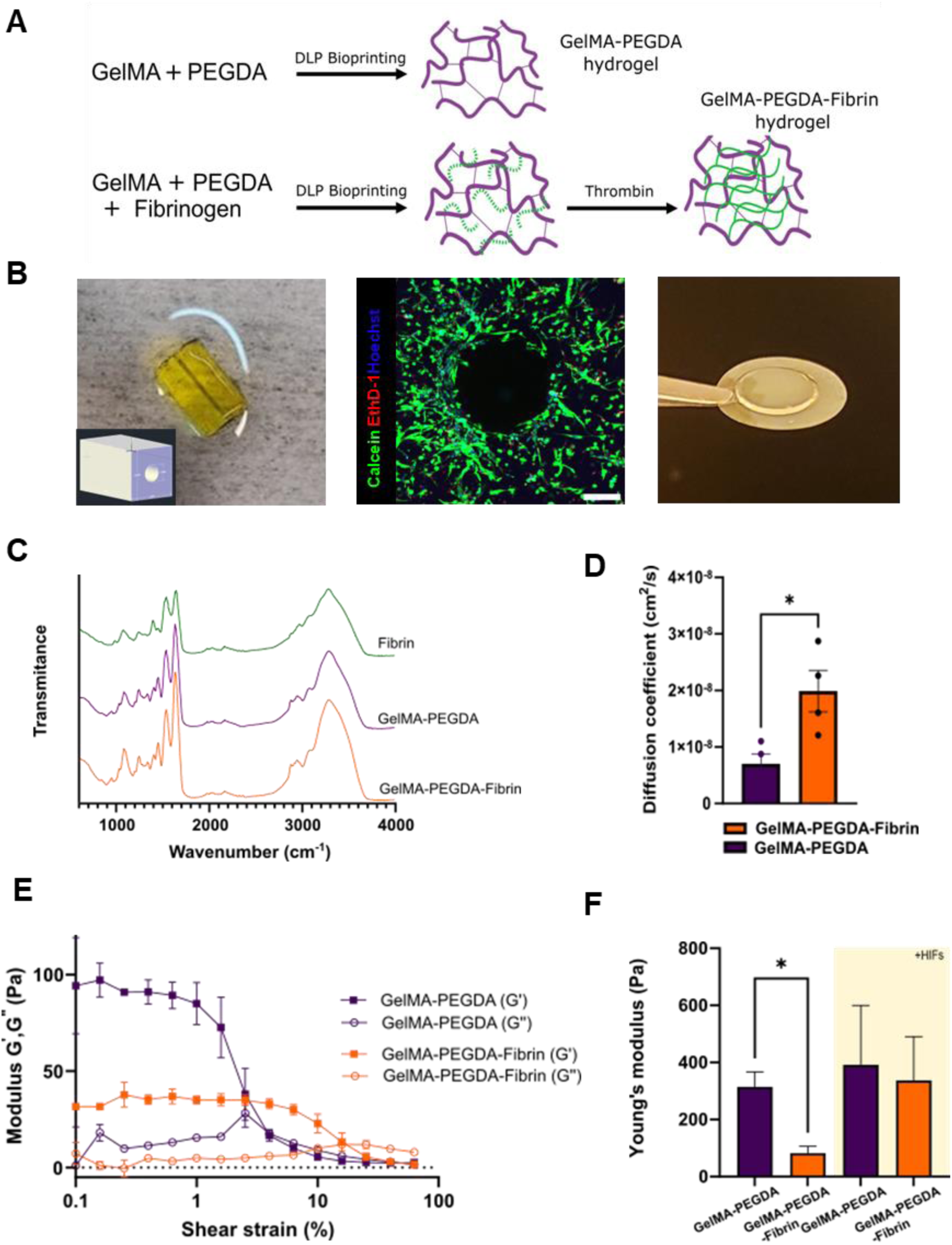
Physico-chemical characterization of bioprinted hydrogels. (A) Schematic illustration of fibrin interpenetrating polymer network (IPN) formation within GelMA-PEGDA hydrogel after thrombin reaction. (B) A CAD model of the designed hollow cuboid structure (3 × 3 × 5 mm) is bioprinted using the LumenX DLP bioprinter, demonstrating the physical integrity and fidelity of the hollow structure (left panel). Maximum-intensity projection of the confocal image shows high cellular viability after 3 days (live cells labeled in green, dead cells labeled in red, and nuclei labeled in blue) (Scale bar: 200 µm) (middle panel). SOLUS 3D bioprinted disc-shaped hydrogel on coverslip (right panel). (C) Chemical characterization of hydrogels using ATR-FTIR analysis. (E) Diffusion coefficient of the bioprinted hydrogels using FD 150 dextran molecule. Results are shown as mean ± SD (min *n* = 3), * *p* <0.05. (E) Rheological curves showing storage modulus (G’) and loss modulus (G”) as a function of shear strain. (F) Younǵs modulus comparison between cell-laden and cell-free hydrogels. Results are shown as mean ± SD (min *n* = 3), * *p* <0.05.

A customized digital light processing stereolithography (DLP-SLA) 3D bioprinting system built from a low-cost Solus 3D printer (Junction3D) ^23^ was used to print disc shape designs with 6 mm in diameter and 0.3 mm in height. The working prepolymer solution was added into the vat preheated to 37 °C. The CAD design was then printed layer by layer onto either silanized 12-mm glass coverslips or Tracketch^®^ polyethylene terephthalate (PET) 10-mm membranes with a pore size of 0.4 μm (it4ip). Once the printing process was completed, the hydrogels were rinsed with warm phosphate-buffered saline (PBS) at 37°C to eliminate any remaining unreacted polymer. Subsequently, they were transferred to the cell culture well plates and/or Transwell^®^ inserts for further experimentation. The samples that included fibrinogen in the bioink composition were incubated in cell culture medium containing 5U/mL thrombin (Merck, Life Science), for 4 h to guarantee a complete reaction between fibrinogen and thrombin, leading to the formation of an interpenetrating network of fibrin within the GelMA-PEGDA hydrogel ^24^.

To assess the print versatility of the hydrogel formulation to fabricate more complex structures, we designed a hollow cuboid structure with outer dimensions of 3 × 3 × 5 mm and a cylindrical central channel of different diameters ranging from 0,3 to 1 mm, resembling a vessel-like geometry. We used the bioink formulation of 5% GelMA, 1.25% PEGDA, 0.2% LAP, and 0.05% tartrazine to print the structure with the LumenX™ DLP bioprinter (Cellink, Sweden). The CAD model was sliced and fabricated under optimized light exposure parameters (18 seconds exposure time with a 4X exposure factor) to achieve high-resolution hollow constructs. Following printing, the samples were gently detached from the platform and rinsed with phosphate-buffered saline (PBS).

### 2.3. Hydrogel characterization

#### 2.3.1. Chemical characterization using Fourier Transform Infrared spectroscopy (FTIR)

The chemical composition of the printed hydrogels was assessed using Fourier-transform infrared spectroscopy (FTIR) with a Nicolet iS 10 instrument from ThermoFisher Scientific. An attenuated total reflectance (ATR) diamond and a DTGS detector were utilized. Hydrogel discs with a diameter of 8 mm and a height of 1 mm were prepared as described before. Moreover, a control sample of fibrin hydrogel was prepared by mixing 200 µl of fibrinogen (0.25% w/v) and thrombin (5 U/mL), which was further incubated at 37°C for 4 h to ensure the reaction completion. Afterwards, all samples were rinsed with Milli-Q water and allowed to dry in a vacuum desiccator for 24 h before the FTIR analysis. Scans were conducted in the range of 500-4000 cm^−1^ with a resolution of 4 cm^−1^. The reported spectra represent an average of 16 scans.

#### 2.3.2. Evaluation of hydrogel mesh size and diffusivity

Since the Flory-Rehner theory is not applicable to estimate the mesh size of hydrogel co-networks ^25^, a diffusion assay was performed by passing dextran fluorescent molecules with different molecular weights through the hydrogels to estimate the cut off size of the pores. For this purpose, 6 mm diameter disc shape designs of both GelMA-PEGDA and GelMA-PEGDA-Fibrin hydrogels were printed onto PET membranes and mounted on Transwell^®^ inserts using double-sided pressure-sensitive adhesive (PSA) rings (Adhesive research) as described in ^26^. They were then placed in standard cell culture conditions overnight, with a temperature of 37°C and 5% CO_2_, to allow for equilibrium swelling.

Fluorescently labeled dextran solutions of MW 4 kDa, 70 kDa, 150 kDa, 500 kDa, and 2000 kDa (Sigma-Aldrich) were individually prepared at a concentration of 0.5 mg/mL in PBS. Subsequently, 200 μL of the pre-warmed dextran solutions were added to the upper compartment (donor) of the Transwell inserts, while 600 μL of PBS was added to the lower compartment (receiver). The plates were incubated at 37°C, and at specified time intervals, 50 μL of liquid was removed from the receiver compartment and replaced with warmed PBS. The collected samples were then transferred to black 96-well plates, and the fluorescence of FITC or Rhodamine was measured using an Infinite M200 PRO Multimode microplate reader (Tecan) at excitation/emission wavelengths of 490/525 nm and 540/625 nm, respectively. The concentrations of dextrans at different time points were estimated using standard calibration curves, and the apparent diffusion coefficients (D) were calculated using the equations (1) and

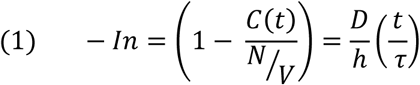

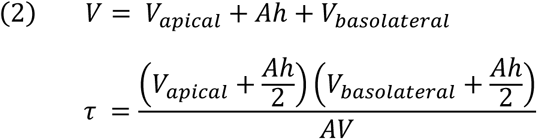

Where C(t) represents the concentration of dextran in the basolateral chamber as a function of time (t). N denotes the total mass of dextran, h represents the height of the hydrogels, V represents the total volume of the hydrogel, and A represents the area where the diffusion takes place.

#### 2.3.3. Mechanical properties

Rheological measurements were performed to assess the mechanical properties of the bioprinted hydrogels. First, hydrogels were bioprinted into 8 mm diameter discs, with and without the encapsulation of human intestinal fibroblasts (HIFs) as stromal cells. Samples with stromal cells were immersed in culture medium, while samples without cells were submerged in PBS for 5 days at 37°C to reach swelling equilibrium. As the sample dimensions were changed due to the swelling, they were punched to achieve 8 mm diameter to test. Rheological measurements were carried out using a Discovery HR − 2 rheometer with parallel sandblasted plates of 8 mm diameter and a Peltier temperature control system. Amplitude sweeps were performed at shear strain amplitude from 0.1% to 1000% and constant 1 rad/s frequency. The temperature during the measurements was maintained at 37°C. Storage (G’) and loss (G”) moduli were measured as a function of strain to determine the value of the modules in the linear viscoelastic (LVE) range, from which the elastic component of the moduli, E, was calculated based on a Poisson’s ratio of 0.5.

### 2.4. Cell culture

Human intestinal fibroblasts (HIFs) primary cells (passage 3 to 8) were kindly provided by Prof. Bruno Sarmento (i3S, Portugal) and were cultured in fibroblast medium (FM) supplemented with 2% fetal bovine serum (FBS), 1% of Fibroblast Growth Supplement (FGS) and 1% of penicillin/streptomycin solution (all from ScienCell).

EndoGRO^TM^ human umbilical vein endothelial cells (HUVECs) were purchased from Merck and cultured (passage 3 to 5) in EndoGRO complete culture medium, which consists of a basal medium and its supplement kit (EndoGRO™ LS Supplement rhEGF, L-Glutamine (200mM), heparin sulfate, hydrocortisone hemisuccinate, FBS and ascorbic acid).

Human colorectal SW480 and HT29 cells were kindly provided by Dr Lorena Dieguez (International Iberian Nanotechnology Laboratory (INL), Portugal) and were cultured in high glucose DMEM (Gibco, ThermoFisher), supplemented with 10 % FBS (Gibco, ThermoFisher) and 1 % Penicillin-Streptomycin (Sigma-Aldrich).

Each cell type was cultured individually in tissue culture flasks and kept in an incubator at a temperature of 37°C and a CO_2_ concentration of 5% in a humidified environment. Cells were passed weekly, and the medium was replaced every two days.

### 2.5. Colorectal cancer spheroids generation and characterization

The cancer spheroids were generated with Sphericalplate 5D® (Kugelmeiers, Erlenbach, Zurich, Switzerland). Colorectal cancer cells (7.5*10^4^ cells) were seeded in 2 ml culture medium, and the cell suspension was vigorously mixed inside each well to achieve uniform distribution of the cancer cells in the microwells, resulting in spheroids of comparable size. Spheroids were cultured for 3 days before encapsulation in hydrogel.

Spheroid circularity and diameter were quantified using the built-in “Analyze Particles” tool in ImageJ. Initially, all images were converted to 8-bit grayscale, and spheroids were manually isolated and segmented from the background using the Otsu thresholding method within the Adjust Threshold tool. After thresholding, images were transformed into binary masks. These binary images were analyzed using the Analyze Particles function, with parameters set to detect objects ranging from 500 pixels² to Infinity in size and 0.00 to 1.00 in circularity. The options Display Results, Exclude on Edges, and Display Outlines were enabled during analysis. For each spheroid, area and perimeter were measured, and circularity was computed using the formula:

Circularity = 4π × (Area) / (Perimeter²)

A value of 1.0 corresponds to a perfect circle, whereas values closer to 0 indicate greater shape irregularity or elongation. Spheroid diameter was estimated by averaging the major and minor axes of the fitted ellipse for each particle. All data were exported from ImageJ and subjected to statistical analysis to evaluate morphological differences across experimental groups.

The viability of spheroids was assessed using a calcein-AM/ethidium homodimer Live/Dead kit (Invitrogen) after 3 days. Spheroids were incubated for 30 minutes with calcein-AM (1 mM; live cells in green) and ethidium homodimer 1 (6 mM; dead cells in red) and then washed with PBS (Sigma-Aldrich, USA). To identify living nuclei, Hoechst staining was used. A confocal laser scanning microscope (LSM 800, Zeiss) was used for imaging.

### 2.6. Fabrication and characterization of 3D CRC models

Cell-laden hydrogels were bioprinted encapsulating HIFs to mimic stromal cells in the TME and cancer spheroids as a tumor compartment. The colorectal cancer spheroids derived from HT 29 or SW480 cell lines were collected after 3 days, washed with PBS and together with HIFs at a cell density of 7.5×10^6^ cells/ml suspended in the prepolymer solutions GelMA-PEGDA or GelMA-PEGDA-Fibrin. The bioinks were promptly transferred to the preheated vat, and the printing process was initiated immediately. To ensure that the cell suspension remained evenly distributed throughout the vat during printing, the prepolymer solution was gently stirred by pipetting. Disc-shaped samples with a diameter of 6 mm and a height of 0.3 mm were printed on silanized PET membranes. Following the printing process, the samples were rinsed with warm cell culture medium to eliminate any residual polymer. Subsequently, the cell-laden bioprinted hydrogels were attached to Transwell® inserts using PSA rings. The samples were then cultured for 8 days in an incubator at 37°C and 5% CO_2_, with the medium being replaced every other day.

HIFs viability inside the 3D printed hydrogel discs was evaluated using a calcein-AM/ethidium homodimer Live/Dead kit (Invitrogen) at 2, 5, and 8 days after printing. Hoechst staining was utilized to identify alive nuclei. Imaging was performed using a confocal laser scanning microscope (LSM 800, Zeiss), and image processing and cell viability quantification were performed using the manual cell counter plugin in Fiji software (available at https://hpc.nih.gov/apps/Fiji, NIH).

The migration of HIFs inside the hydrogel bulk was monitored by time-lapse videos using a Leica® THUNDER imaging system under 5% CO_2_ at 37°C. Images were captured every 5 minutes for 10 h and analyzed using the MTrackJ plugin in ImageJ software to track migration of the cells. Data analysis was performed using a custom-made code in Matlab (Mathworks, USA). Cell centroid positions (x,y,z) during the experiment were defined as **r**_i_ = **r**(iDt), being Dt the time between consecutive images and **r** a vector. The vector difference between the initial (t = t_0_) and the final point (t = tf) was defined as the displacement vector **d** and its module ||**d**|| as the net displacement. Furthermore, directionality (||**d**||/||**r**||) and velocity (||**r**||/ (t_f_-t_0_)) were evaluated for each cell.

To assess spheroid growth and morphology, brightfield images were acquired at multiple time points (days 3, 5, and 7) and analyzed using ImageJ software. Spheroid diameter and circularity were measured as previously described. To determine the growth rate, the change in diameter over time was calculated relative to the initial diameter measured after one day in hydrogel. The percentage increase in diameter was calculated using the following formula:

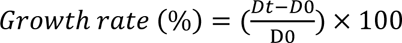

Where Dt is the spheroid diameter at a given time point, and D0 is the initial diameter.

### 2.7. Immunofluorescence analysis

The morphology and phenotype of the spheroids before encapsulation and after embedding in hydrogel in combination with HIFs was characterized by immunofluorescence. Samples were fixed with 10 % neutral buffered formalin solution (Sigma-Aldrich), permeabilized with 0.5% Triton X-100 (Sigma-Aldrich) and blocked with a blocking buffer containing 1% BSA (Sigma-Aldrich), 3% donkey serum (Millipore), and 0.2% Triton X-100 in PBS for at least 2 h at room temperature (RT). Then, they were incubated overnight at 4°C with the corresponding primary antibodies. Then, the samples were incubated with secondary antibodies for 2h at RT. The list with all the antibodies and the dilutions used can be found in the supplementary data. After immunostaining, the samples were mounted on thin glass coverslips using Fluoromount G (Southern Biotech) and imaged using a confocal laser-scanning microscope (LSM 800, Zeiss).

### 2.8. Statistical analysis

The graphs were plotted using the GraphPad Prism 8 version. The data are presented in the figures as the mean ± standard deviation (SD). The number of experiments is reported in each figure. Statistical comparisons were performed using ANOVA test and p-values < 0.05 were considered to be significant.

## 3. Results and discussion

### 3.1. The bioprinted hydrogels reproduce the physico-chemical properties of CRC tumor microenvironment

GelMA-PEGDA hydrogels have been broadly used in the biofabrication of different tissue models due to their combination of cell compatibility, high tunability of mechanical properties, and the possibility to encapsulate cells ^16^. This hydrogel composite has also shown great potential as ink for bioprinting. Recently, we have optimized a GelMA-PEGDA bioink with low polymer content for the bioprinting of cell-laden intestinal scaffolds ^29^. This bioink containing 5% w/w GelMA, 3% PEGDA, 0.4% LAP and 0.025% tartrazine showed very good printability of high-aspect ratio villi-like structures while maintaining the mechanical properties characteristic of the intestinal tissue (around 2 kPa). Additionally, we embedded stromal fibroblasts that reproduced the epithelial-mesenchymal crosstalk. In this work, we aimed to further decrease the PEGDA and LAP content in order to obtain a softer hydrogel that mimics better the mechanical properties of CRC tissue (in the range of 0.1-1 kPa^17^). Thus, we evaluated the printability and functional relevance of the bioink containing 5% GelMA, 1.25% PEGDA, 0.2% LAP and 0.025% tartrazine using two different DLP-based printers: a customized SOLUS 3D printer for printing disc-shaped hydrogels, and the Cellink LumenX bioprinter for printing vessel-like structures (Figure 1B). In this latter case, in order to obtain an open channel along the 5 mm long structure, we increased the tartrazine concentration to 0.05% w/v to better confine the photopolymerization reaction and reduce overexposure in the central channel. The successful fabrication of this complex geometry highlights the high resolution and structural fidelity of our formulation, supporting its suitability for advanced tissue modeling applications. Moreover, the hydrogel supported the encapsulation of HIFs, which exhibited high cell viability after 3 days, confirming the biocompatibility of the hydrogel. To facilitate physicochemical and cellular characterization and ensure consistent experiments, we decided to use the Solus 3D bioprinter to fabricate simple disc-shaped hydrogel samples, that offer a more practical and reproducible approach for performing in vitro assays with an easier handling and uniform cell distribution.

Also, to increase RGD motifs in hydrogel composition and provide better integrin binding, we explore the incorporation of 0.25% fibrinogen in the bioink to obtain a fibrin IPN entangled within the GelMA-PEGDA structure. Fibrin hydrogels have been shown to mimic 3D environment of the tissue, supporting cell adhesion and migration ^30^. However, bioprinted GelMA-PEGDA-Fibrin IPN hydrogels have not been explored yet. We evaluated the effect of the fibrin incorporation on the physicochemical properties of the hydrogel. First, we qualitatively analyzed the hydrogel’s chemical bonds using ATR-FTIR. However, fibrin and GelMA-PEGDA hydrogels showed a very similar IR spectra, so the IPN GelMA-PEGDA-fibrin did not show new significant transmittance bands (Figure 1C). Fibrin and gelatin are natural polypeptides that have many amide bonds (C=O-N-H) in their amino acid chains.

The incorporation of the fibrin hydrogel entangled within the GelMA-PEGDA hydrogel mesh could affect the diffusion properties of the resulting IPN hydrogel. In 3D *in vitro* models, the matrices must allow for the exchange of nutrients, oxygen, and the removal of waste products. The porous structure of the bioprinted hydrogels influences these mass transfer properties. To better understand this, the diffusion profile of the FD 150 kDa dextran compound with 17 nm molecular size was studied, and its apparent diffusion coefficient through the hydrogel co-networks was measured (Figure 1D). We observed that the diffusion coefficient (D) of the dextran through GelMA-PEGDA-Fibrin hydrogel (0.7 ± 0.3×10^−8^ cm/s^2^) is significantly lower than GelMA-PEGDA hydrogel (2 ± 0.7×10^−8^ cm/s^2^), suggesting that successful incorporation of the fibrin IPN in the GelMA-PEGDA hydrogel, affecting the resulting hydrogel mesh size.

Another critical factor influencing cellular behavior within a 3D matrix is stiffness. Cancer spheroids respond to mechanical cues of the surrounding microenvironment, which affect their growth, morphology, and interaction with stromal cells ^31^. To evaluate the stiffness of both hydrogel compositions, we measure the mechanical behavior by rheometry. The elements of the complex shear modulus (G’ and G’’) were measured as a function of shear stress. The graphs showed a characteristic behavior of viscoelastic solids (G’ > G” at rest) with a well-defined linear viscoelastic (LVE) range. The incorporation of the fibrin secondary network did not increase storage modulus, as reported for other IPNs ^32–34^, but resulted in a softer and more elastic hydrogel (Figure 1E). This may indicate that the fibrin could affect the degree of crosslinking of the GelMA-PEGDA hydrogel network, decreasing the value of the storage module and increasing the LVE range. To better measure the tissue mechanical characteristics of the tumor stromal microenvironment, hydrogels were loaded with HIFs and cultured for 5 days before performing the rheological measurements. Cell-laden hydrogels showed higher Young’s modulus compared to cell-free hydrogels. In particular, for the GelMA-PEGDA-fibrin IPN hydrogel, the stiffness increased from 82 ± 42 Pa to 337 ± 268 Pa (Figure 1F), which accounts for the presence of the stromal HIF cells and the secretion of ECM ^35,36^. The stiffness values of the bioprinted hydrogels fall within the range characteristic of CRC tissue (0.1-1 kPa^17,18^), confirming that they effectively reproduce key aspects of the native TME. Click or tap here to enter text.

### 3.2. The presence of the fibrin network within the GelMA-PEGDA hydrogel promoted cell elongation and migration of embedded human intestinal fibroblasts

Fibroblasts are considered one of the predominant stromal cells in almost all tissues ^37^. Thus, we encapsulated human intestinal fibroblasts (HIFs) to mimic the stromal microenvironment of the CRC. For this, HIFs were mixed with prepolymer solution of GelMA-PEGDA or GelMA-PEGDA-Fibrin and bioprinted in hydrogel discs. Cell viability after printing was evaluated using a live/dead assay and confocal microscopy. Two days after encapsulation, the cells were uniformly distributed within the hydrogel and started to elongate as observed with the Live/Dead assay (Figure 2A). Overall, cell viability remained above 90% during the initial week post-encapsulation in both hydrogels, regardless of the fibrin content (Figure 2B).

**Figure 2.**
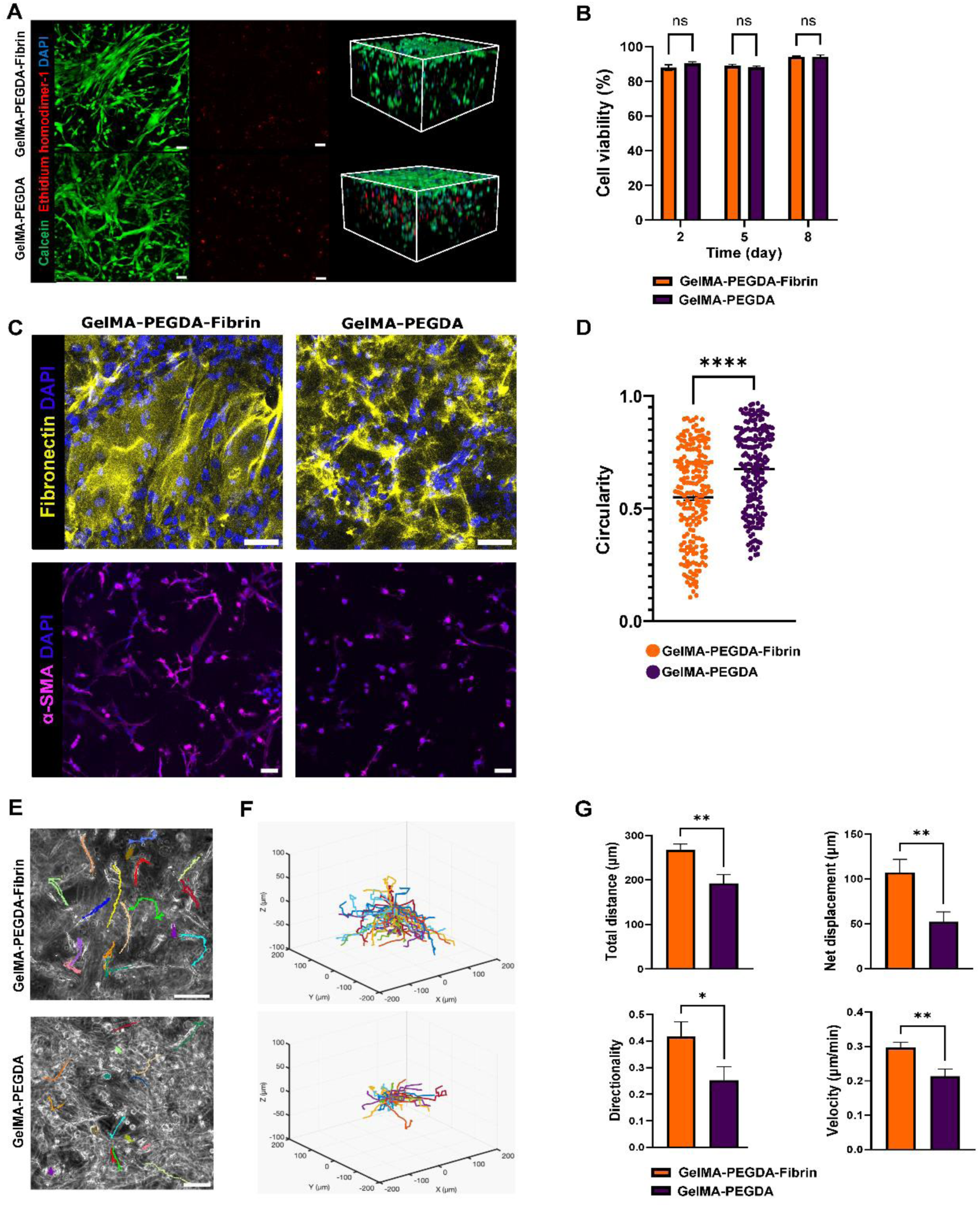
Stromal cell viability and migratory behavior inside the bioprinted hydrogels. (A) Live/dead assay maximum projection of confocal images (left panel) and spatial distribution (right panel) of embedded HIFs after two days post-encapsulation (live cells labeled in green, dead cells labeled in red, and nuclei labeled in blue) (Scale bar: 50 µm). (B) Quantification of cellular viability at different time points after encapsulation. (C) Maximum intensity projections of HIFs encapsulated in GelMA–PEGDA (left) and GelMA–PEGDA–Fibrin (right), immunostained for fibronectin (yellow) and α-smooth muscle actin (α-SMA, purple) markers (Scale bar: 50 µm). (D) Cell morphology quantification of the circularity of embedded HIFs after 5 days. Results are shown as mean ± SD (min *n* = 3), **** *p* < 0.0001. (E) Representative final frame images of the time-lapse video after cell tracking analysis (Scale bar: 100 µm). (F) Representative plots of the cell’s trajectory which were analyzed by MATLAB analyzing software. (G)Quantification of the cell’s trajectory for different aspects of cell migration including total distance (µm), net displacement (µm), directionality, and velocity (µm/min). Results are shown as mean ± SD (min *n* = 3), ** *p* < 0.005; * *p* <0.05.

Fibrin plays a key role in regulating fibroblast behavior by providing structural and mechanical cues that promote cell elongation and alignment. Fibroblasts adhere to and reorganize fibrin fibers, generating traction forces that realign the matrix and guide their orientation^38^. As shown in Figure 2C, immunofluorescence characterization for fibronectin marker demonstrated clear differences in ECM deposition. In GelMA–PEGDA–Fibrin hydrogels, fibronectin was deposited in elongated and aligned structures, indicating that fibroblasts adopted an oriented morphology and organized their extracellular matrix accordingly. In contrast, HIFs within GelMA–PEGDA hydrogels showed a more random distribution of fibronectin, with a disorganized network lacking a preferred orientation. To further evaluate HIFs elongation in both hydrogel compositions, immunofluorescence for α-smooth muscle actin (α-SMA) marker was performed after 5 days of encapsulating fibroblasts. Fibroblasts displayed a more elongated morphology in the GelMA-PEGDA-Fibrin hydrogel bulk compared to the GelMA-PEGDA hydrogel. The differences in cell shape were quantified by measuring the circularity of fibroblasts in the hydrogel bulk, where values near 1 indicate a more rounded shape, and lower values represent increased elongation. The data show a statistically significant decrease in circularity for fibroblasts in the GelMA–PEGDA–Fibrin group compared to those in the GelMA–PEGDA group, indicating enhanced cell spreading (Figure 2D).

Cell migration is one of the fundamental phenomena in cancer metastasis ^39^. To assess how the hydrogel composition might affect cell migration, we recorded time-lapse videos three days after encapsulating fibroblasts within the hydrogels (Figure S1). We monitored the movement of HIFs by manually tracking their paths within the hydrogel for 10 hours (Figure 2E) and then plotted the normalized individual trajectories to evaluate the randomness of movement (Figure 2F). Qualitatively, cells migrating within hydrogels containing fibrin displayed longer trajectories, a characteristic not limited to the xy plane but extending into the z-axis as well. This suggests that fibrin facilitates the migration of cells in all directions, with similar extension observed in all three dimensions. Quantitatively, the trajectories of cells migrating in fibrin-containing hydrogels differed from those in hydrogels without fibrin (Figure 2G). Cells covered a greater total distance (267 ± 50 µm compared to 192 ± 74 µm), resulting in higher velocities (0.296 ± 0.055 µm in GelMA-PEGDA-Fibrin versus 0.213 ± 0.082 µm in GelMA-PEGDA). Additionally, the net displacement traveled by cells in GelMA-PEGDA-Fibrin (107 ± 54 µm) exceeded that in GelMA-PEGDA alone (52 ± 43 µm). Remarkably, this also led to enhanced directionality, with cells migrating in fibrin-containing hydrogels exhibiting twice the directional behavior compared to those in GelMA-PEGDA alone. These results together with the fibronectin staining suggest that fibrin promotes a guided migration through the hydrogel network. Other works have also observed an accelerated invasion, elongation, and migration of fibroblasts within fibrin-based soft hydrogels ^40^.

### 3.3. CRC tumor spheroids maintain proliferation and characteristic phenotype

In order to evaluate the bioprinted hydrogel performance mimicking the CRC tumor microenvironment, spheroids from two CRC cell lines of different phenotype were generated. HT29 cells are characterized by a less invasive, epithelial-like phenotype, whereas SW480 cells have a more invasive, mesenchymal-like phenotype ^41^. Spheroids were generated using Sphericalplate5D and cultured and characterized over a week to evaluate the best time point for hydrogel encapsulation (Figure 3A). At day 3 after generation, both HT29 and SW480 spheroids exhibited high cellular viability (labeled in green) and few dead cells (labeled in red) (Figure 3B).

**Figure 3.**
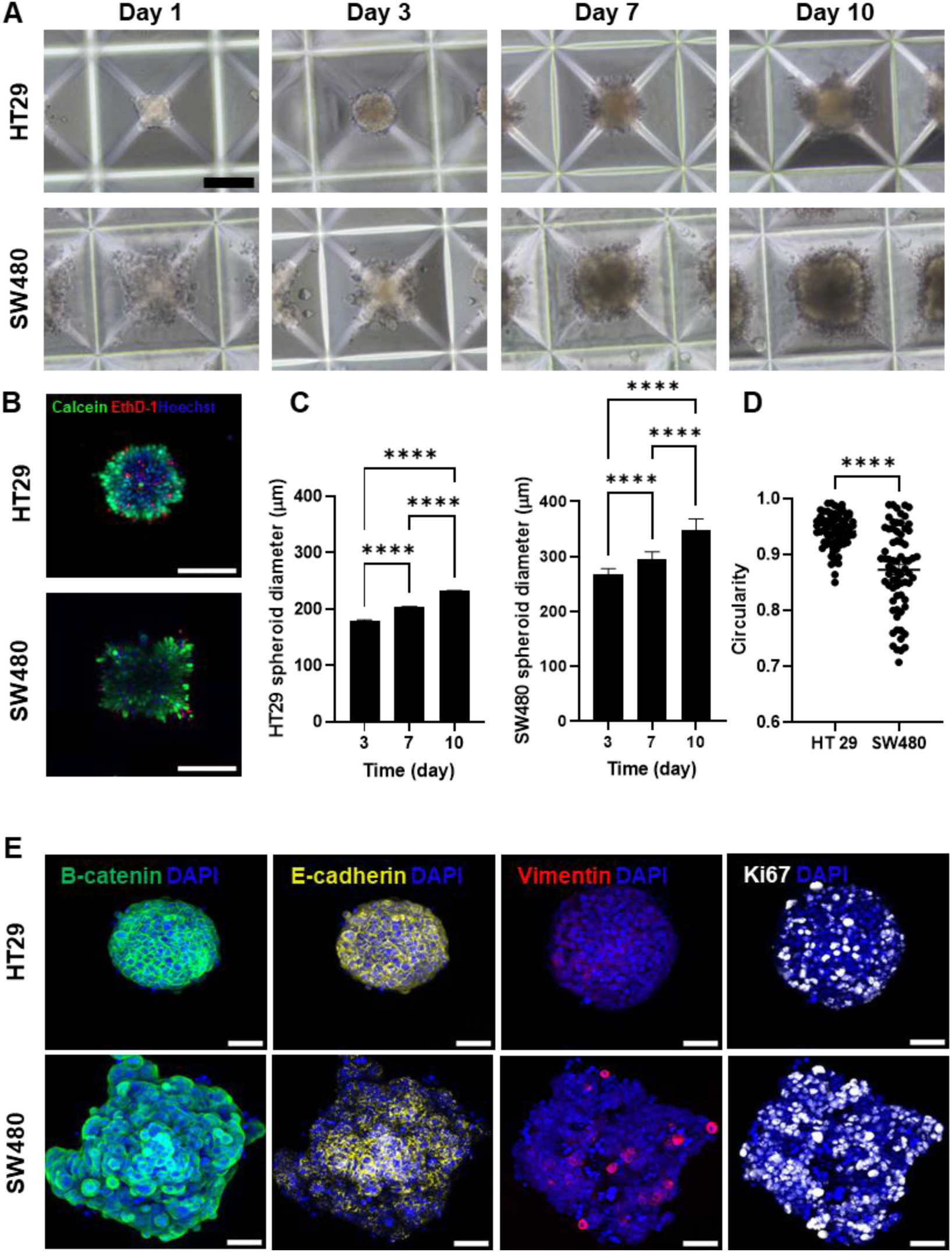
Spheroid characterizations. (A) Brightfield images showing the growth of HT29 and SW480 spheroids over 10 days of culture within Sphericalplate 5D® microwells. Scale bar: 100 µm. (B) Maximum intensity projection of confocal images demonstrates high cellular viability in both HT29 and SW480 spheroids after 3 days. (live cells labeled in green, dead cells labeled in red, and nuclei labeled in blue) (Scale bar: 50 µm). (C) Spheroid diameter quantification for HT29 spheroids (left panel) and SW480 spheroids (right panel) after 3, 7, and 10 days. Results are shown as mean ± SD (n ≥ 60), **** *p* < 0.0001. (D) Spheroid circularity quantification on day 3. Results are shown as mean ± SD (n ≥ 70), **** *p* < 0.0001. (E) Maximum intensity projections of immunostainings for β-catenin (green), E-cadherin (yellow), vimentin (red), and Ki67 (white) markers on HT29 and SW480 spheroids on day 3. Scale bar: 50 µm

Spheroid growth, morphology, and circularity are important factors that indicate the degree of cell-cell adhesion, uniformity of cell distribution, and overall structural integrity. Moreover, it can provide insight into the intrinsic properties of the cancer cell line, such as epithelial vs. mesenchymal phenotypes, proliferative behavior, and metastatic potential ^42^. Even though starting from the same number of cells, SW480 spheroids showed bigger diameter and looser structure after formation, corresponding to the low cell-cell adhesion characteristic of this metastatic cell line. Upon culture, both cell lines spheroids retained cell proliferation showing a gradual and significant increase in diameter, with the SW480 spheroids showing greater expansion, growing from 267 ± 11 µm (day 3) to 347 ± 21 µm (day 10). In comparison, HT29 spheroids displayed a more compact structure with a slower growth trend, reaching from 179 ± 15 µm (day 3) to 232 ± 8 µm (day 10) (Figure 3C). To further investigate the morphological characteristics of HT29 and SW480 spheroids, circularity was quantified after 3 days of spheroid generation. HT29 spheroids maintained high circularity values, ranging from 0.85 to 0.99, reflecting a compact and spherical structure. In contrast, SW480 spheroids displayed notably lower circularity, ranging from 0.70 to 0.98 (n≥70) (Figure 3D), indicating the presence of outward protrusions and cells spreading from the spheroid periphery, correlating with its mesenchymal phenotype ^44^.

Further characterization with immunostaining revealed that 3 days after formation spheroids retained their characteristic phenotypes (Figure 3E). HT29 spheroids showed a compact and rounded morphology with membrane-localized E-cadherin and β-catenin, and negligible vimentin expression characteristic of epithelial primary tumors with high cell-cell adherent junctions^44,46^. In contrast, SW480 spheroids showed lower E-cadherin expression, higher vimentin expression, and more cytoplasmic or nuclear β-catenin localization. Moreover, the Ki67-positive proliferative cancer cells were more abundant and distributed in SW480 spheroids compared to HT29 spheroids, which have been shown to express more in mesenchymal cells ^47^.

### 3.4. GelMA-PEGDA-Fibrin hydrogel network promoted the tumor cell invasion of bioprinted HT29 and SW480 spheroids

3D CRC models were finally obtained by bioprinting GelMA-PEGDA-Fibrin or GelMA-PEGDA hydrogels embedding HIFs stromal cells and HT29 or SW480 spheroids after 3 days formation. Even though the size of the spheroids at the time of encapsulation, spheroids were retained between the bioprinted bioink layers and were evenly distributed inside the hydrogels. HT29 spheroids retained their compact round shape after printing. However, SW480 spheroids, due to their loose cell-cell adhesion, exhibited partial dissociation upon encapsulation, resulting in heterogeneity in spheroid size and morphology across both hydrogel types. The 3D CRC hydrogel models were cultured over 8 days post-encapsulation and the growth and circularity of the spheroids were monitored as key indicators that reflect how the physicochemical and mechanical properties of hydrogels influence tumor behavior, including proliferation, structural integrity, and invasiveness over time ^31,43^. HT29 spheroids embedded in GelMA–PEGDA–Fibrin were larger with more irregular and asymmetric shapes; in contrast, they were more compact and rounded in GelMA–PEGDA hydrogels (Figure 4A). This visual observation was supported by quantitative analysis of spheroid circularity (Figure 4B), which showed a significantly lower circularity value in the GelMA–PEGDA–Fibrin hydrogel (0.88 ± 0.05) compared to the GelMA–PEGDA hydrogel (0.91 ± 0.04). A lower circularity indicates disrupted spheroid boundaries and the presence of protrusive structures, which are characteristic of an invasive phenotype ^31^. These changes were also significant when size and growth rate were quantified. HT29 spheroids embedded in the GelMA–PEGDA–Fibrin hydrogel exhibited a more substantial increase in size, growing from 300 ± 34 µm on day 3 to 342 ± 64 µm by day 7, while those in the GelMA–PEGDA hydrogel expanded from 284 ± 32 µm to 321 ± 46 µm during the same period (Figure 4C). This translated to a significantly higher growth rate up to 39 ± 3% by day 7 in GelMA-PEGDA-Fibrin compared to 31± 2 % in GelMA–PEGDA hydrogel (Figure 4D). These morphological changes suggest that the softer and more permissive fibrin-enriched hydrogel allows cancer cells to spread and reorganize more freely, promoting invasive growth patterns.

**Figure 4.**
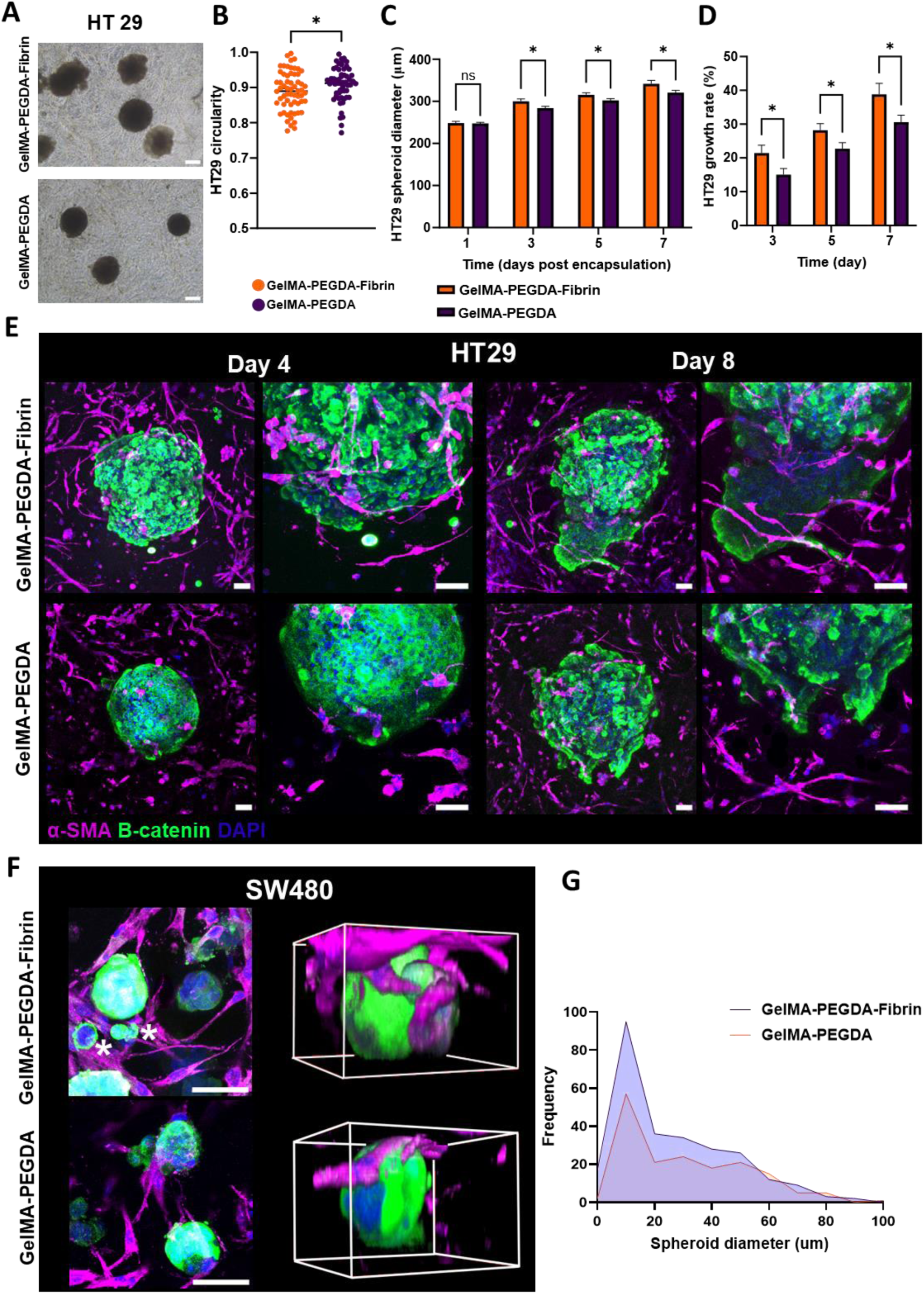
The Effect of hydrogel composition on HT29 and SW480 spheroid morphology and tumor–stroma interactions. (A) Brightfield images of HT29 spheroids in GelMA–PEGDA and GelMA–PEGDA–Fibrin hydrogels after 7 days. Scale bars: 200 μm. (B) Spheroid circularity quantification on day 7. Data are presented as mean ± SD (n ≥ 60); *p < 0.05. (C) Quantification of HT29 spheroid diameter in GelMA–PEGDA and GelMA–PEGDA–Fibrin hydrogels over time (days 3, 5, and 7). Data are presented as mean ± SD (n ≥ 35); ns (not significant), *p < 0.05. (D) Relative spheroid growth rate in GelMA–PEGDA and GelMA–PEGDA–Fibrin hydrogels over time (days 3, 5, and 7) calculated with respect to initial diameter after one day. Data are presented as mean ± SD (n ≥ 35); *p < 0.05. (E) Immunostaining for β-catenin (green) and α-SMA (magenta) markers on HT29 spheroids co-cultured with human intestinal fibroblasts (HIFs) after 4 and 8 days in GelMA–PEGDA and GelMA–PEGDA–Fibrin hydrogels. DAPI was used to stain the nuclei. Scale bars: 50 μm. (F)Maximum intensity projections of immunostainings for α-SMA (magenta) and β-catenin (green) on SW480 spheroids co-cultured with human intestinal fibroblasts (HIFs) after 8 days in GelMA–PEGDA–Fibrin and GelMA–PEGDA hydrogels. The spatial distribution of spheroids and fibroblasts interactions in the hydrogel bulk is shown in a 3D reconstruction (right panels). DAPI was used to stain the nuclei. Scale bars: 50 μm. (G) Quantitative analysis of spheroid diameter distribution in GelMA-PEGDA-Fibrin and GelMA-PEGDA hydrogels. Data are presented as mean ± SD (n ≥ 10).

HT29 spheroids inside the CRC 3D hydrogel were further characterized by immunofluorescence, revealing notable differences in spheroid morphology and fibroblast interaction (Figure 4E). After 4 days in the GelMA–PEGDA–Fibrin hydrogels, the HT29 spheroids expressing β-catenin began to show signs of invasion, with less compact morphology and an irregular border, while α-SMA-positive fibroblasts started to spread and interact with them. In comparison, within GelMA-PEGDA hydrogel, the HT29 spheroids retained a compact structure with minimal invasion into the surrounding stroma, and fibroblasts remained more rounded with limited interaction. By day 8, the differences became more obvious. In GelMA–PEGDA–Fibrin, HT29 spheroids showed highly irregular borders and invasive morphology, suggesting enhanced interaction with the surrounding stromal microenvironment. This was accompanied by improved fibroblast elongation and interaction with spheroids. Meanwhile, in GelMA–PEGDA, spheroids mostly retained their spherical shape with smooth edges, and fibroblasts remained relatively rounded and less interactive, indicating limited tumor-stromal interactions.

In the case of SW480 tumor model, the heterogeneity of spheroid size due to the partial disruption of the spheroids during encapsulation impaired the systematic quantification of size and growth rate (Figure S2). However, a clear difference was observed between the two types of hydrogels in the immunostaining images (Figure 4F). In GelMA–PEGDA–Fibrin hydrogels, SW480 spheroids showed the dissociation of individual tumor cells that migrated away from the spheroid (demonstrated with asterisks) and interacted with surrounding HIFs. Moreover, HIFs exhibited elongated morphologies that were closely interacting with both spheroid and dissociated tumor cells, suggesting that these elongated stromal cells may facilitate tumor cell dissemination by creating supportive migratory pathways. By contrast, spheroids in GelMA–PEGDA remained more intact, with fewer cells escaping from the spheroid periphery and limited stromal integration. The 3D reconstruction views (right panels) supported these findings by showing dissociated single tumor cells migrating from SW480 spheroids and interacting with elongated HIFs in the fibrin-based hydrogel. However, SW480 spheroids within GelMA–PEGDA retained a compact structure, with no apparent single-cell dissemination, and fibroblasts had a less elongated morphology. The 3D rotational projection highlighted these differences more clearly on day 8 (Figure S3). Quantitative analysis of spheroid diameter distribution further supported these findings (Figure 4G). In GelMA–PEGDA–Fibrin, a higher frequency of spheroids with diameters below 20 µm was observed, consistent with increased dissociation of tumor cells from the spheroid. By comparison, smaller spheroids and single cells were less frequent in GelMA–PEGDA, indicating reduced levels of spheroid dissociation and tumor cell escape.

Overall, the invasive behaviors of HT29 and SW480 spheroids in GelMA–PEGDA–Fibrin versus GelMA–PEGDA hydrogels highlight how hydrogel composition and mechanical properties influence tumor invasiveness and fibroblast behavior. We specifically selected two colorectal cancer cell lines with distinct phenotypes to capture different modes of invasion. HT29 cells exhibit a more epithelial phenotype, so their invasiveness was primarily reflected in spheroid growth, morphological changes, and reductions in circularity. In contrast, SW480 cells display a mesenchymal-like phenotype with weaker cell–cell adhesion, and their invasiveness was more evident through single-cell dissociation and migration into the surrounding stroma. In both cases, the softer, fibrin-enriched hydrogels promoted enhanced invasiveness. The incorporation of fibrin reduced hydrogel stiffness, creating a permissive environment that facilitates cell spreading and migration. Notably, fibroblasts embedded in the fibrin-enriched hydrogel became more elongated and aligned, forming organized interactions with spheroids and dissociated cancer cells, which likely assisted tumor cell migration ^44^. Softer matrices also allow greater responsiveness to enzymatic degradation and ECM reorganization, which are critical for tumor cells to navigate the extracellular matrix and form invasion pathways ^45^. These results demonstrate that hydrogel mechanical properties can modulate invasion across different tumor phenotypes, supporting the use of tunable fibrin-enriched hydrogels as a versatile bioprinted platform for studying colorectal cancer progression.

## Supporting information

Suppplemental video S3A

## Acknowledgments

The project leading to these results has received funding from “la Caixa” Foundation (ID 100010434), under the agreement LCF/PR/HR20/52400011, and Grant PID2021-129115OB-I00 funded by MCIN/AEI/ 10.13039/501100011033 and by “ERDF A way of making Europe”. MPK thanks the Severo Ochoa program of the Spanish Ministry of Science and Competitiveness for a predoctoral grant (PRE2020-092849). The IBEC group was supported by the department of Research and Universities (2021 SGR 01495) and the CERCA Programme of the Generalitat de Catalunya. The authors wish to acknowledge the MicroFabSpace and Microscopy Characterization Facility, Unit 6 of ICTS “NANBIOSIS” from CIBER-BBN at IBEC.

