## Supplemental Table S1 for "Engineering a soft tumor microenvironment: fibrin-enriched hydrogels promote cancer cell invasion in 3D bioprinted colorectal cancer models"

**Supplementary information**

Table 1. The list of antibodies

**Primary antibodies**

Anti- β catenin (1:200, Abcam, ab2365)

Anti- E cadherin (1:1500, BD biosciences 610181)

Anti- Vimentin (1:100, Santa Cruz, sc6260)

Anti-Ki67 (1:100, BD Pharmingen 550609)

Anti-α-smooth muscle actin (α-SMA, 1:200, Abcam, ab7817)

Anti-Fibronectin (1:100, Sigma, F7387)

**Secondary antibodies**

Anti-mouse Alexa Fluor 647 (1:500, Invitrogen, A-31571)

Anti-rabbit Alexa Fluor 568 (1:500, Invitrogen, A-10042)

Anti-goat Alexa Flour 488 (1:500, Invitrogen, A-11055)

**Nuclei staining**

DAPI (1:1000, Thermo Fisher D1306)
