## Supplemental Figure S2 for "Engineering a soft tumor microenvironment: fibrin-enriched hydrogels promote cancer cell invasion in 3D bioprinted colorectal cancer models"

**Supplementary information**

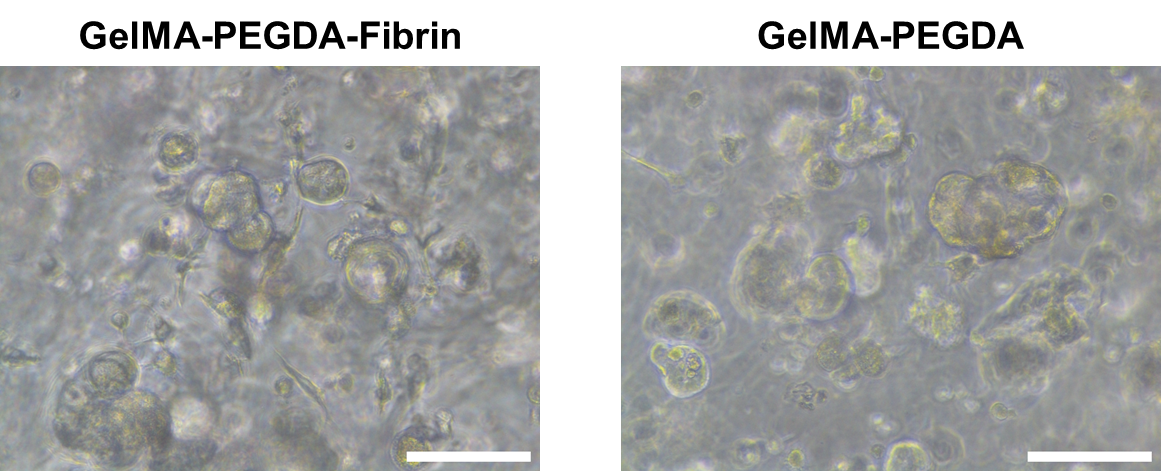

Figure S2. Representative brightfield images of SW480 spheroids in GelMA–PEGDA and GelMA–PEGDA–Fibrin hydrogels after 7 days of culture. Scale bars: 100 μm.
